# Statistically principled feature selection for single cell transcriptomics

**DOI:** 10.1101/2024.10.11.617709

**Authors:** Emmanuel Dollinger, Kai Silkwood, Scott Atwood, Qing Nie, Arthur D. Lander

**Affiliations:** Center for Complex Biological Systems, University of California, Irvine, Irvine, CA 92697; Department of Developmental and Cell Biology, University of California, Irvine, Irvine, CA 92697; Department of Mathematics, University of California, Irvine, Irvine, CA 92697

## Abstract

The high dimensionality of data in single cell transcriptomics (scRNAseq) requires investigators to choose subsets of genes (feature selection) for downstream analysis (e.g., unsupervised cell clustering). The evaluation of different approaches to feature selection is hampered by the fact that, as we show here, the performance of feature selection methods varies greatly with the task being performed. For routine cell type identification, even randomly chosen features can perform well, but for cell type differences that are subtle, both number of features and selection strategy can matter strongly. Here we present a simple feature selection method grounded in an analytical model that, without resorting to arbitrary thresholds or user-defined parameters, allows for interpretable delineation of both how many and which features to choose, facilitating identification of biologically meaningful rare cell types. We compare this method to default methods in scanpy and Seurat, as well as SCTransform, showing how greater accuracy can often be achieved with surprisingly few, well-chosen features.

## INTRODUCTION

Single cell RNA sequencing (scRNAseq) measures the transcriptional profiles of individual cells, enabling cell type classification, lineage inference, and elucidation of experimental differences in both gene expression and cell type abundance (Das et al., 2022; Junttila et al., 2022; Nguyen et al., 2021; Simmons, 2022; Tam & Ho, 2020; Tritschler et al., 2019; Xie et al., 2021). As cell type identity is generally not known *a priori*, it is traditional to use unsupervised clustering to group transcriptomically similar cells. As a first step, genes (“features”) considered most likely to be markers of cell type or state are selected and used for subsequent dimensionality reduction and clustering (Heumos et al., 2023).

In general, limiting features to those that are most informative for downstream applications improves interpretability, increases computational efficiency, prevents overfitting, and improves the performance of clustering algorithms (Li et al., 2017). However, there is no one-size-fits-all definition of “most informative”. Most widely-used algorithms calculate gene expression variation across cells, but they differ significantly in how variation is measured, as well as how the appropriate number of features is determined (Andrews & Hemberg, 2018; Brennecke et al., 2013; Hafemeister & Satija, 2019; Jiang et al., 2016; Lall et al., 2022; Satija et al., 2015; Stuart et al., 2019; Su et al., 2021; Townes et al., 2019; Tyler et al., 2024). Although some comparisons have been made of the effects of different feature selection methods on clustering (Germain et al., 2020; Su et al., 2021; Zhao et al., 2024), it occurred to us that performance of a selection method will likely depend on aspects of the dataset to which it is applied. These could include how many cell types are present, their relative abundance, and the number and magnitude of gene expression differences between cells of different types. To our knowledge, the interaction of these factors with different methods for ranking and selecting genes has not been explored in a systematic way.

Here we show that, for datasets in which the desired goal is to cluster relatively abundant cells that differ greatly in gene expression, how features are selected is almost irrelevant. Even random sets of genes, if large enough, tend to perform nearly as well as algorithmically-chosen features. In more demanding situations, in which random feature selection does not suffice for clustering, we identify cases in which both the method of feature selection and the number of features used markedly influence clustering outcomes.

This led us to revisit the question of how best to identify genes for which cell-to-cell variation is greater than otherwise expected. Starting with an analytical model for the structure of un-normalized scRNAseq data, we developed a method for quantifying both degree of expression variation and probability of occurrence by chance. This method, which we call BigSur (<u>B</u>asic <u>I</u>nformatics and <u>G</u>ene <u>S</u>tatistics from <u>U</u>nnormalized <u>R</u>eads), provides a theoretical framework for scRNAseq data analysis which enables both feature selection and the inference of gene regulatory networks from gene-gene correlations (the latter use outlined elsewhere (Silkwood et al., 2024)). Here we show that using BigSur for feature selection enables identification of biologically relevant groups of cells while minimizing loss of discriminatory power due to the use of excessive numbers of features.

## RESULTS

### For common tasks, random sets of genes can perform as well as algorithmically-chosen features

Existing feature selection algorithms often choose up to several thousand genes for use in cell clustering. As this can represent a substantial fraction of the expressed genome, one is left to wonder how many of these features are necessary, and whether the number chosen is particularly important.

In common scenarios, in which cell types of interest are relatively abundant and well separated in gene expression space, we find that features chosen at random often perform nearly as well as those selected by popular algorithms. We illustrate this by clustering the 10k cell PBMC (peripheral blood mononuclear cell) dataset from 10x Genomics, which is commonly used for methods evaluation. Initially, we used scanpy’s default feature selection method (highly variable genes, “HVGs”), with cutoffs and parameters set to their default values. HVGs bins genes by mean expression and ranks each gene by their z-score calculated from the genes in that bin (Satija et al., 2015). Using the 3643 genes chosen by HVGs, cells were clustered according to the default pipeline: finding the first 50 principal components, creating a nearest neighbor graph in PCA space, and clustering using the Leiden algorithm. Clusters were then labeled based on the expression of known marker genes. As expected, cell groups were well separated on a UMAP plot (Figure 1A, left; see also Supplementary Figure 1).

**Figure 1:**
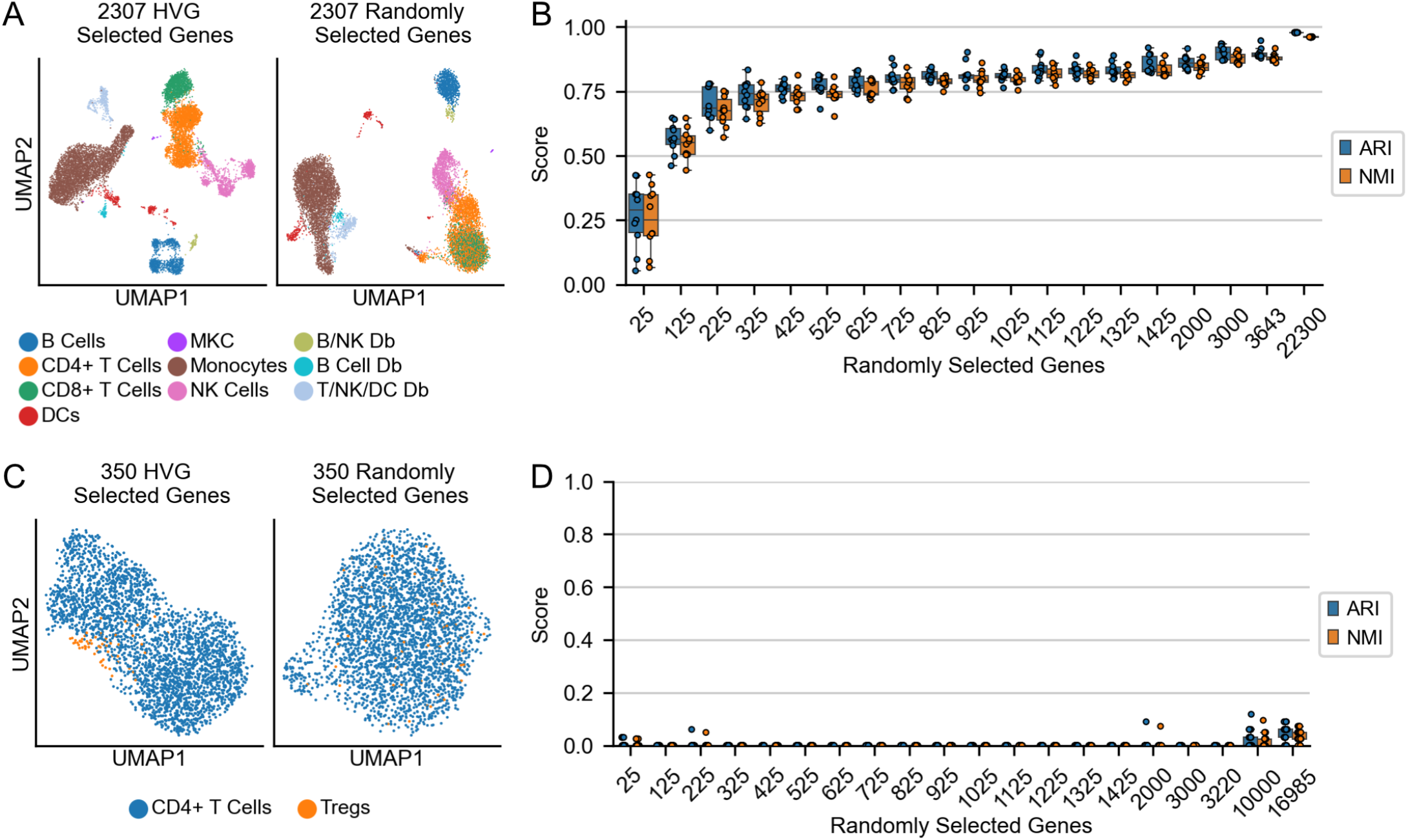
Performance of randomly selected genes as features. **A**. UMAP representation of PBMCs calculated using either HVG-selected genes (left) or the same number of randomly selected genes (right). **B**. Adjusted Rand index (ARI) and normalized mutual information (NMI) of cell type identification using a Support Vector Machine (SVM) on the PCs, as a function of number of genes selected. **C**. UMAP of CD4+T cells, including the T regulatory cell (Tregs) subset, calculated using either HVG-selected genes (left) or same number of randomly selected genes (right). **D**. ARI and NMI of Treg identification, as a function of number of randomly selected genes.

For comparison, we used an equal number of genes (3643) chosen at random as features, and performed the same task. As shown in Fig. 1A, right, the same cell types grouped well and separated similarly from each other, with the one exception that some of the CD4+ and CD8+ T cells were intermixed.

We next asked how many random genes were needed to cluster cells correctly, and how repeatable the results were. Since labels are already assigned to each cell in this case, we can treat this question as a supervised problem. We therefore set as ground truth the cell type labels identified using HVGs (Fig. 1A) and assessed the ability of a straightforward classifier (a linear support vector machine, or SVM) to identify the cell types from the 50 top PCs of randomly selected genes (see methods). We varied the number of genes from 25 to 22,300, testing 10 samplings at each size, retraining the SVM at each step.

As the numbers of randomly selected genes increased, both the adjusted Rand index (ARI) and the normalized mutual information (NMI)—two common classification metrics— increased quickly to an ARI of 0.8 at 725 genes and an NMI of 0.8 at 925 genes (Fig. 1B). Both scores continued to slightly rise up to 0.98 and 0.96 for ARI and NMI respectively with all genes. The variance (across the 10 trials) of the ARI and NMI scores decreased as the number of genes increased, but was already low by 525 genes (Fig. 1B). These results show that almost any random set of genes of size greater than a few hundred is potentially sufficient to correctly classify groups of cells that are well separated in gene expression space. Although we are not aware of this observation having been explicitly noted before, it supports the view, voiced by others, that the effective dimensionality of gene expression is far lower than the number of expressed genes (Heimberg et al., 2016; Lenz et al., 2016; Lukk et al., 2010). In other words, any random sample of reasonable size would be expected to include a substantial number of genes that correlate with all major patterns of gene expression.

### Some tasks are sensitive to choice of feature selection algorithm

Given the results above, the task in Fig. 1A is clearly not suitable for assessing the performance of any feature selection algorithm, as the lack of an algorithm (using all genes) or a trivial algorithm (random gene selection) perform extremely well.

This led us to consider more challenging tasks, for example one involving more subtle gene expression differences. We thus subsetted just the CD4+ T cells in the PBMC dataset, and subclustered them. Using the 350 features selected by HVGs, we could identify a FOXP3+ T regulatory cell (Treg) cluster of approximately 1.8% (53/2908) of the cells (Fig. 1C, left; Figure S1). Tregs are a well described CD4+ T cell subtype, regulating immune responses in many biological systems (Miragaia et al., 2019). In this case, using an equal number of random genes as features, the Treg population was not identifiable (Fig. 1C, right). Even using much larger numbers of genes—up to all 16,985 significantly expressed genes–and testing 20 random gene sets for each sample size, ARI and NMI scores remained close to zero (Fig. 1D). Thus, in a sufficiently difficult task, HVGs performed much better than random.

### Number of features can influence success or failure in unpredictable ways

The fact that random gene selection failed to identify Treg cells even when the entire expressed transcriptome was used emphasizes that a good feature selection must not only include enough genes that are predictors of cell types or states, but also exclude enough genes that are not. This reflects a well known phenomenon in machine learning, in which the accuracy of a classifier initially increases with increasing number of features and then decreases (Hughes, 1968). This happens because, in high dimensions, consideration of data that have low predictive power can “swamp” out the effects of data with high predictive power (i.e., the “noise” can overwhelm the “signal”) (Zimek, 2012).

Because of this, it is important that feature selection algorithms for scRNAseq not only order genes by their utility as features, but also decide how far down the list one should go, i.e. the number of features to use, when performing unsupervised clustering. Currently popular algorithms typically choose feature number based either on arbitrary variability cutoffs or on calculations that depend upon arbitrarily adjustable hyperparameters (Hafemeister & Satija, 2019; Hao et al., 2021; Wolf et al., 2018).

To assess how important these choices are, we selected three clustering tasks: subclustering CD4+ T cells (as in Fig. 1C-D); CD8+ T cells (also from the PBMC 10K data set), and human retinal amacrine cells (clustering and cell type identification is shown in Figure S2) (Menon et al., 2019). In each case, we varied the number of features used, starting from those considered by HVGs to be most highly variable to the least. We calculated the fraction of the cell state of interest in each resulting cluster, and report the fraction of the “best” cluster (the cluster with the most cells of the cell state of interest), which we refer to as the “purity score” (see methods).

As shown in Figure 2, the qualitative results differed markedly among the three tasks. With amacrine cells, a population of SLC12A7-expressing cells could be cleanly identified with either 150 or 3424 features, the latter representing HVG’s default selection (Fig. 2A). In this case, purity was relatively stable regardless of the number of features used (Fig. 2B).

**Figure 2:**
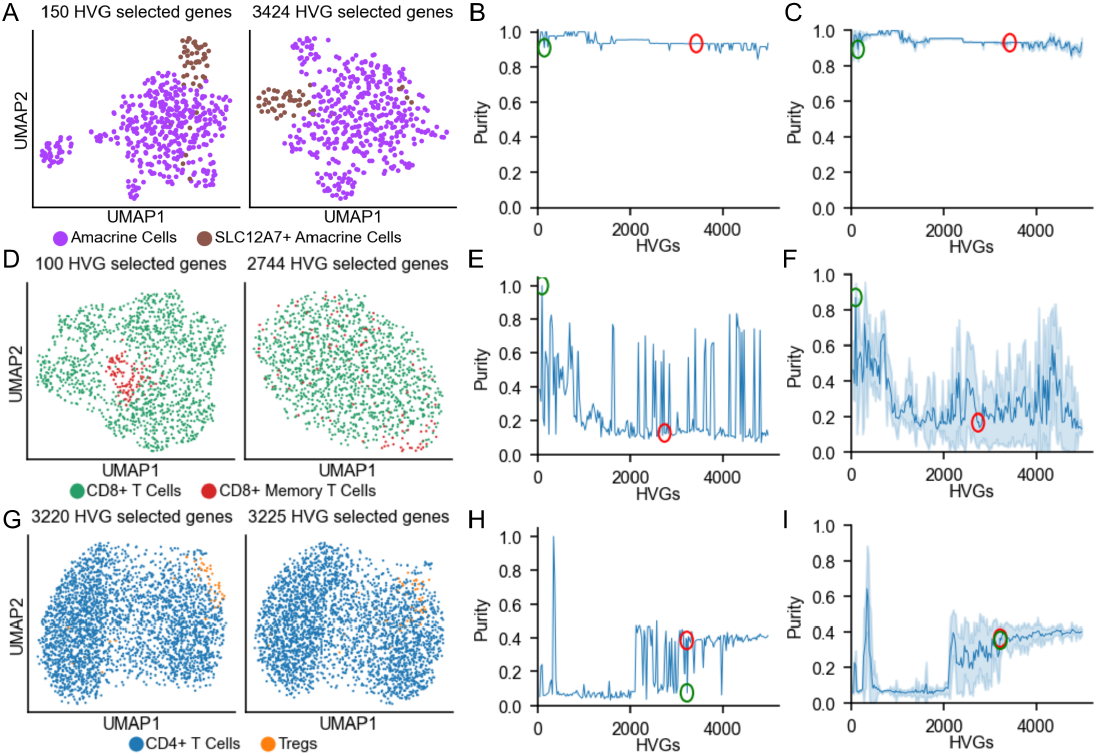
Numbers of features and feature selection method influence clustering. **A**. UMAPs of amacrine cells and SLC12A7-expressing amacrine cells using either 150 HVGs (left) or the default number of HVGs (3424; right). **B**. Purity score for SLC12A7+ cells, in clusters derived from amacrine cells using different thresholds for numbers of HVG features. Green and red circles show results using 150 and 3424 genes, respectively. **C**. Mean and standard deviation of purity scores for SLC12A7+ cells, using 50 randomly selected starting seeds for the Leiden clustering algorithm. **D**. UMAPs of CD8+ T cells and memory CD8+ T cells using 100 HVGs (left) and the default number of HVGs (right). **E**. Purity score as the number of HVGs increases. Green and red circles mark enrichment scores at 100 HVGs and the default number of HVGs (2744), respectively. **F**. Mean and standard deviation of purity scores using 50 different Leiden starting seeds, as in C. **G**. UMAPs of CD4+ T cells and Regulatory T cells (Tregs) using either 3220 genes (left) and default number of HVGs (right, 3225). **H**. Purity score of Tregs as in panels B and E. **I**. Mean and standard deviation purity scores of 50 randomly selected Leiden starting seeds, as in panels C and F.

With CD8+ T cells, we identified a subpopulation of memory T cells (a CCL5+ population equaling 9.3% of the total) using the top 100 HVG features (Fig. 2D), but performance degraded markedly when larger numbers of features were used, and sometimes declined to very low levels (Fig. 2E). Importantly, using the default number of features suggested by HVGs (2744; red circle in Fig. 2E), the ability to identify memory T cells as a distinct cluster was lost.

Finally, we turned again to the CD4+ T cells. The Treg population previously discussed (Fig. 1C) mostly grouped together when the default number of 3220 HVGs was used (Fig 2G-H, Fig. 2H red circle), but adding in even 5 more features destroyed the ability to identify this cluster (Fig. 2H, green circle).

It is worth commenting that the observed “choppiness” in panels 2E and 2H—where purity scores jump from high to low with the addition or subtraction of just a few features—partly reflects the sensitivity of Leiden clustering to the choice of random starting seed (Traag et al., 2019). Accordingly, we also carried out the analyses in panels 2B, E and H using 50 randomly selected starting seeds at each feature number, and plot the mean and standard deviation of the results in panels C, F and I, respectively.

Taken together, the results in Fig. 2 indicate that, for tasks in which having an algorithm for feature selection matters, selecting the right number of features can be important, affecting both the average and the range of purity scores one could expect to encounter in practice. Too many features can be as harmful as too few, and within some ranges, even small changes in feature number can have dramatic consequences. Importantly, feature numbers chosen by commonly used procedures under default conditions would be expected to lead to systematic misclassification. We therefore asked whether there might be a principled way to optimze the joint tasks of finding features and determining how many to use.

### A statistical approach to feature selection

Common feature selection methods tend to choose genes by the degree to which they are variable within a dataset, with differences among methods typically involving the way data are initially transformed, the way variability is defined and measured, and how the appropriate number of features is determined. Ideally, one would want to select features that are more variable than expected by chance, but this requires knowledge of the null distribution, i.e. the data distribution to expect when all cells are biologically equivalent (i.e., the only sources of variability being technical and biological noise). Because there has long been a lack of general agreement on the appropriate null distribution for scRNAseq data, it has become common to estimate one empirically from the data, by fitting gene expression data to some kind of flexible model (e.g. negative binomial). This is, for example, the principle used by SCTransform, which also takes advantage of such fitting to normalize gene expression values (Hafemeister & Satija, 2019; Satija et al., 2015; Stuart et al., 2019).

Defining “null” using data in which cell type differences are expected to exist is fundamentally problematic, but might be justified if the genes that differ between cell types are relatively few in number. Unfortunately, the results in Fig. 1 suggest this is rarely a good assumption. The fact that relatively small numbers of random genes can often be used as features to drive correct cell clustering tells us that gene expression differences associated with cell types are, in fact, pervasive, involving a large fraction of the genome.

We therefore chose the alternative approach of constructing a null distribution analytically, based on assumptions about how scRNAseq data arise. Specifically, we assume that biological noise—random fluctuations in transcript numbers in the same or identical cells—follows a log-normal distribution, which agrees with both theoretical predictions and empirical observations in mammalian cells (Bahar Halpern et al., 2015; Beal, 2017). Beyond this, we assume that the technical noise associated with cell preparation, library preparation, sequencing, etc., can simply be approximated as sampling noise, i.e., by the Poisson Distribution. This agrees with a number of recent observations and arguments in the scRNAseq literature (Choi et al., 2020; Kim et al., 2015; Sarkar & Stephens, 2021; Wang et al., 2018).

We thus write the following model:

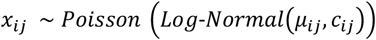

where μ_*ij*_ is the expected number of transcripts of gene *j* in cell *i*, and *c*_*ij*_ is the coefficient of variation of that value, and *x*_*ij*_ is the observed counts (UMI) detected for gene *j* in cell *i*. Note that, in our terminology, we parametrize the log-normal distribution in terms of its actual mean and coefficient of variation, and not the mean and coefficient of variation of the underlying normal distribution from which it may be derived.

To guide the use of this model in developing statistics for the analysis of real-world data, we first generated synthetic data potentially representative of a situation in which two cell types or states are present in a dataset, and differ only in the expression of a known number of genes (Figure 3). Specifically, we generated data for 2,000 cells and 15,000 genes expressed at a range of levels, using an underlying coefficient of variation of gene expression of 0.7 for each gene (Fig. 3A; see Methods). For a fraction of genes, the two cell types—which are present in equal proportions—were made “truly variable”, i.e. they drew from distributions with different means; for the rest of the genome the cells drew from a single distribution for each gene. For simplicity, in Fig. 3 we do not include additional effects due to differential sequencing depths among cells (we discuss this issue later on), and values were chosen independently for every gene.

**Figure 3:**
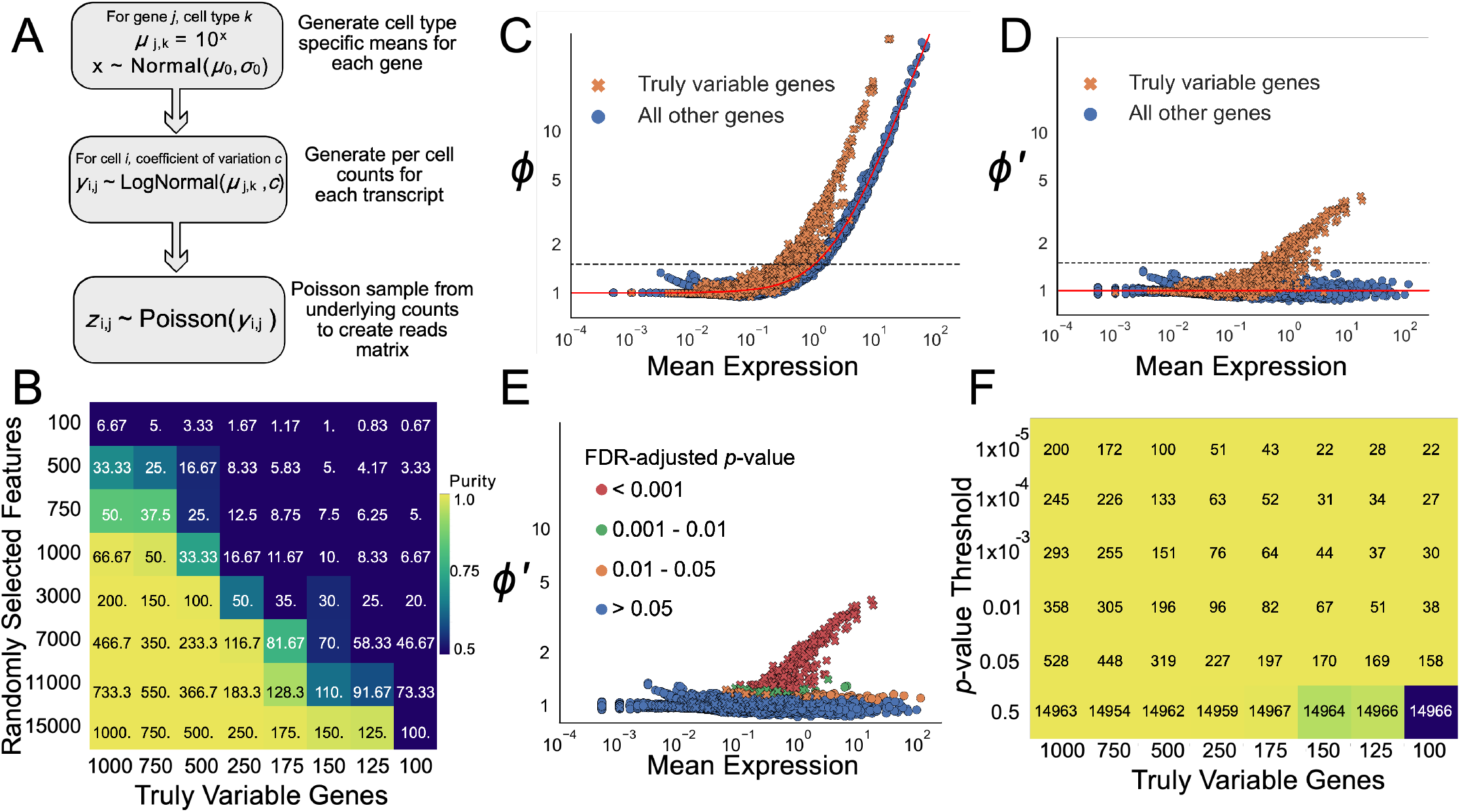
BigSur identifies truly variable features. **A**. Synthetic data generation pipeline (see Methods). **B**. Average purity of clusters (see Methods) generated using genes selected at random as features. Each purity score is the average of ten trials of random feature selection. The numbers on each tile indicate the expected number of truly variable genes to be found among the selected features. **C**. Observed relationship between the Fano factor and the mean for simulated data. Orange crosses indicate genes that were truly variable, i.e., were selected from different distributions for the two different cell types. Blue dots are genes that were selected from the same distributions for the two cell types. The red line depicts the expected relationship, under the null hypothesis, between the Fano factor and mean expression. The dashed line at *ϕ* = 1.5 shows the large number of non-truly-variable genes that would be chosen if the Fano factor is used to select features. **D**. Observed relationship between the modified corrected Fano factor (*ϕ’*) and mean expression. The red line again depicts the expected relationship under the null hypothesis. Using *ϕ’* as a feature criterion avoids the inclusion of non-truly-variable genes. **E**. Points from panel D colored by their FDR-adjusted *p*-value. Markers are as in panels C-D. **F**. Purity scores (color scale as in panel B) for clusters obtained using features selected by BigSur at different *p*-value thresholds. Underlying data were identical to those in panel B. Numbers overlaid on each tile indicate the number of features selected by BigSur in each case.

If we generate a large number of datasets in this manner, in which the number of truly variable genes varies from 100 to 1000 we immediately see that there is a threshold above which randomly selected features (genes) begin to work very well in correctly classifying the two cell types. The threshold depends both on how many features are truly variable and how many random genes are selected as features, but tends to occur when the number of truly variable genes expected to be found among the random features lies in the vicinity of 30-150. Interestingly, the larger the set of random features one uses, the greater the fraction of truly variable features they must include in order to perform well; this demonstrates how both signal (inclusion of truly variable features) and noise (exclusion of non-truly variable features) both matter for classifying high dimensional data (Zimek et al., 2012).

Ideally, an algorithm for identifying features should seek to compare the observed variance of a feature with the variance expected under the null hypothesis, i.e., when features are not truly variable. For data that are Poisson-distributed, variance is equal to the mean, so the ratio of observed variance to mean, also known as the Fano Factor, equals 1 under the null hypothesis. Hereinafter we use *ϕ*_*j*_ to stand for the Fano factor of gene *j*, 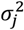 the observed variance of gene *j*, and *μ*_*j*_ the observed mean. Thus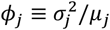.

Fig. 3C shows the distribution of observed Fano factors for synthetic data generated as in Fig. 3B, in which the number of truly variable genes was set to 1,000. Although one should expect to observe *ϕ*>1 for truly variable genes, many non-variable genes also display large values of *ϕ*; in particular this is true for highly expressed genes. This reflects the fact that the variance of a Poisson sample from any distribution is equal to the variance of the Poisson distribution (i.e., the mean) plus the variance of that distribution. As we may equivalently express variance as the square of the coefficient of variation times the mean, i.e., (*cμ*)^2^, a modified Fano factor expressed as:

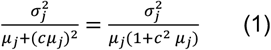

should have an expectation value of 1 for a compound Poisson distribution. Indeed, using this metric on the data in Fig. 3C, one no longer observes large values for highly expressed genes, and non-variable genes are cleanly separated from truly variable ones (Fig. 3D).

Although the modified Fano factor serves as an appropriate metric for identifying truly variable genes in the synthetic data in Fig. 3, it is not suitable for use on real-world single cell RNA sequencing data without an additional correction. The reason is that, in real-world data, cells are typically sequenced to widely different depths. Normalizing gene expression to read depth recovers an estimate of gene expression in each cell, but the distribution of that estimate will not be the same as if each cell had actually been sequenced to the same depth. Accordingly, the expectation value under the null hypothesis for expression (2) will not be 1. As recently pointed out (Lause et al., 2021), one way to correct for this is not to scale gene expression values, but instead correct their Pearson residuals. The Pearson residual, a measure of deviation from the mean, is defined, in cell *i* and gene *j*, as

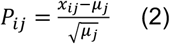

where *μ*_*j*_ is the mean expression over all cells. The Fano factor may then be expressed as an average of squared Pearson residuals:

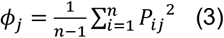

Lause et al. noted that, when sequencing depth is variable, the expectation value for *ϕ*_*j*_ can be made equal to 1 if the term *μ*_*j*_ in the Pearson residual of every cell *i* is replaced by *μ*_*ij*_, an estimate of the mean specifically for that cell (i.e. an estimate of the global mean scaled by the total sequencing depth of the cell) (Lause et al., 2021). If we further modify that term by dividing by 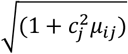, we obtain 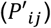, which we refer to as a “modified corrected Pearson residual”:

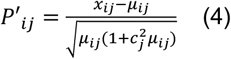

as well as a “modified corrected Fano factor”, 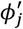 :

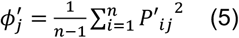

Note that *P*′_*ij*_ and 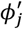 are calculated from raw, unnormalized counts, as it is the *μ*_*ij*_, and not the *x*_*ij*_, that undergo correction for sequencing depth variation.

Because we may expect 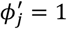 under the null hypothesis, observing 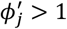 should identify features likely to be truly variable. To do so in a principled way, however, requires knowing the probability of observing any value of 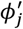 under the null hypothesis. While it is generally not possible to obtain an analytical expression for the full distribution of 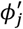, we may construct an arbitrary number of moments of that distribution from the moments of the distributions of the individual *P*′_*ij*_, which we can in turn construct from the moments of the Poisson-Log Normal distribution parametrized by *μ*_*ij*_ and *c* in each cell (Silkwood et al., 2024). Given a sufficient number of moments, procedures exist for estimating tail probability densities (Cornish & Fisher, 1938). In this manner, one can associate any value of 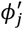 with a *p*-value, i.e. the probability of observing it by chance. Given a set of *p*-values, one can further identify a *p*-value threshold for any level of false discovery (Benjamini & Hochberg, 1995). In Fig. 3E, the data in panels C-D are colored by the false discovery rate (FDR) threshold interval into which their *p*-values fall. It is immediately apparent that, for these data, few non-variable genes display FDR-adjusted p-values less than 0.05, and none display values less than 0.001.

We have named the algorithm for jointly computing modified corrected Fano factors and their *p*-values *BigSur*, which stands for <u>B</u>asic <u>I</u>nformatics and <u>G</u>ene <u>S</u>tatistics from <u>U</u>nnormalized <u>R</u>eads. Fig. 3F applies BigSur to all eight datasets (columns) in Fig. 3B, reporting both the number of features that satisfy different significance thresholds, and their success in clustering the two cell types (the latter represented using the same color scheme as in Fig. 3B). A surprisingly small number of features suffice to produce good clustering. For example, in the dataset containing only 100 truly variable genes, as few as 22 statistically significant features recovered pure clusters, whereas no number of random features (from 100 to 15,000) could do so. This reflects the ability of BigSur to recover a large fraction of true positives while minimizing false discovery.

The only hyperparameter used by BigSur is *c*_*j*_, the underlying coefficient of variation of gene expression (i.e. the actual biological variation among equivalent cells). While this number may, in principle, differ for each gene, experimental studies suggest it does not vary greatly (Bahar Halpern et al., 2015), and we provide a method here to estimate a consensus value of *c*_*j*_ (which we hereafter simply call *c*) from a full gene expression dataset (see Methods).

It is worth noting that the modification and correction of Pearson residuals may also be generalized to higher order statistics, such as correlation coefficients, and that this can be leveraged to more accurately assess gene-gene correlations (Silkwood et al., 2024).

### BigSur helps identify rare, biologically relevant cell states

We next examined the performance of BigSur on experimental data, focusing on cases in which the presence of rare cell types, or subtly different cell types, might be expected to pose challenges for clustering. First, we chose the CD4+ T cell subset from the 10k PBMC set. As shown in Figure 4A-B, the Fano factors associated with gene expression rose sharply with expression level, similar to what was observed with synthetic data (Fig. 3C). In contrast, the modified corrected Fano factors (using a fitted *c* = 0.25) were generally mean-independent, with most genes displaying a value of *ϕ*′ near 1. These results suggest that the statistical properties of real data resemble those of the synthetic data; they also suggest that, in this dataset, the patterns of expression of most genes are consistent with not being truly variable. For a small subset of genes, however, values of *ϕ*′ up 10 were observed, and most of those above 1.5 were associated with FDR-adjusted *p*-values less than 0.05. Overall, 156 genes were characterized by *ϕ*′ >2 and *p* <0.05 (Fig. 4C). Among these was *FOXP3*, considered a marker for regulatory T cells (Treg).

**Figure 4:**
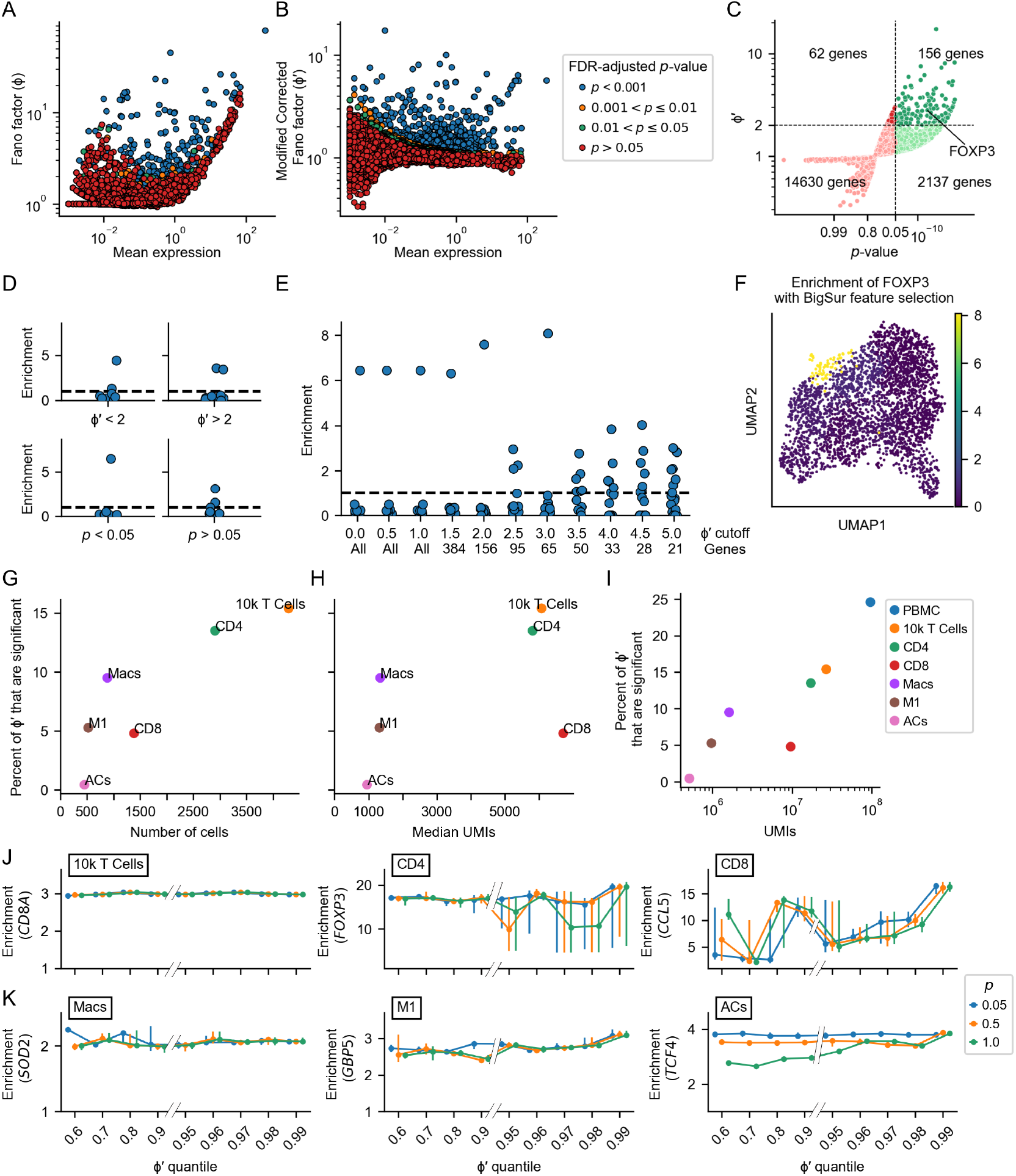
Clustering performance using values and significance levels of modified corrected Fano factors. The Fano factor *ϕ* (**A**) and the modified corrected Fano factor *ϕ’* (**B**) were calculated for all genes in the CD4+ T cell dataset, and are plotted as a function of gene expression level, and colored according to the *p*-value of *ϕ’*. **C**. For the same dataset, values of *ϕ’* and their *p*-values are plotted against one another. Green and red distinguish *p*-values smaller or greater than 0.05, respectively. Bold and pale colors distinguish values of *ϕ’* greater or smaller than 2, respectively. **D-E**. Enrichment for *FOXP3* after Leiden clustering using features determined by different *p*-value and *ϕ’* cutoffs. Enrichment was defined as mean expression of *FOXP3* within each cluster divided by mean *FOXP3* expression across the dataset. Each point represents an individual cluster. **D**. Enrichment scores using solely a *ϕ’* threshold (top) or solely an adjusted *p*-value threshold. **E**. Using an FDR-adjusted *p*-value threshold of 0.05, the *ϕ’* threshold was progressively increased from 0 to 5 (note drop in number of selected genes). **F**. UMAP of the CD4+ T cell dataset with *ϕ’* cutoff of 3 and p-value cutoff of 0.05. Cells are colored by their cluster’s enrichment. **G-H**. For each of six datasets, the percentage of *ϕ’* that were significant (FDR < 0.05) was plotted against the number of cells (**G**) and the median sequencing depth (**H**) (see main text for information on the datasets). **I**. For the datasets in panel F, as well as the full PBMC dataset (Fig. 1), the percentage of *ϕ’* that are significant was plotted against the total number of transcripts in the entire dataset (which reflects both sequencing depth and number of cells). Note how different datasets vary greatly in both total amount of sequence and evidence for cellular heterogeneity (percentage of *ϕ’* that are significant). **J-K**. For the datasets in panel F, features were selected using different quantile thresholds for *ϕ’* (abscissa) and different *p*-value cutoffs (by color). The Leiden clustering algorithm was run on the selected features using 40 different randomly chosen starting seeds. The enrichment of a gene marking a particular group of cells of interest was calculated for each of the resulting clusters: *CD8A* for the 10k T Cells dataset, *FOXP3* for the CD4 dataset, *CCL5* for the CD8 dataset, *SOD2* for the Macrophage dataset (Macs), *GBP5* for the M1 macrophage dataset and *TCF4* for the amacrine cells dataset (ACs). For each Leiden seed, the enrichment level in the cluster with the greatest enrichment was found. Each point on the plot indicates the median of this value over the different Leiden seed choices, and the bars show the inter-quartile range (25% to 75% of the data).

To investigate which factor, *ϕ*′ or the *p-*value, played a greater role in enabling identification of Tregs as a unique cluster, we carried out Leiden clustering using features identified at different thresholds for these parameters, calculating a *FOXP3*-enrichment score for each cluster produced. The enrichment score was calculated by dividing the mean normalized expression of *FOXP3* in each cluster by the mean expression of *FOXP3* in the dataset. As shown in Fig. 4D-E, use of an adjusted *p*-value threshold of 0.05 was particularly important to achieve good cell separation, i.e. the inclusion of non-statistically significant features appears to be especially detrimental to rare cell type identification. Among statistically significant features, the magnitude of *ϕ*′ appeared to be less important, unless the cutoff became so high that the absolute number of selected features became very small—somewhere in the vicinity of 50-150 genes (Fig. 4E).

We next extended this analysis to five other datasets that contained known cell “subtypes”: the full T cell subset of the 10k PBMC dataset, the CD4 and CD8 subsets of those T cells, a macrophage dataset (Carvalho et al., 2021), an M1 macrophage subset of those, and the retinal amacrine cell subset. These datasets span a wide range of at least three characteristics: the fraction of all *ϕ*′ observed to be statistically significant (*p*<0.05), the number of cells sequenced (Fig. 4G), and the median sequencing depth (median UMI/cell) (Fig. 4H). The latter two of these may be expected to have a strong influence on the statistical power to identify differences—fewer cells mean fewer observations, and lower sequencing depth means more observations that are zero and thus minimally informative. An alternative metric that captures a sense of the combined statistical power of these two observations is the total number of UMI in the entire dataset (i.e. the product of the number of cells and the mean UMI/cell). Plotting this against the fraction of *ϕ*′ values that is statistically significant (Fig. 4I) shows that, overall, the significant *ϕ*′ fraction declines with decreasing total UMI. This general pattern is to be expected, as loss of statistical power should correlate with identification of fewer *ϕ*′ as significant, however the magnitude of the effect varies with dataset (for example, that fraction is particularly low for CD8 T cells). Such variation provides a measure of cellular heterogeneity—i.e. when the fraction of *ϕ*′ that is significant is unexpectedly low, it suggests that gene expression differences between cells are especially subtle.

We next examined how well Leiden clustering relying on BigSur for feature selection performed in enriching for markers of known cell subsets in each dataset. For example, we used enrichment for *CD8A* to assess clustering of CD8 T cells from mixed T cells; *FOXP3* for Tregs from CD4 T cells; *CCL5* for memory T cells from CD8 T cells, *SOD2* for M1 macrophages from mixed macrophages, *GBP5* for a subset of M1 macrophages, and *TCF4* for glycinergic amacrine cells from total amacrine cells (Figure S3). In each case we considered three different FDR-corrected *p*-value thresholds of 0.05, 0.5 and 1.0 and varied the *ϕ*′ cutoff. To facilitate comparisons between datasets, the *ϕ*′ cutoff was varied by quantile, i.e. 0.6 means the top 40% of *ϕ*′ values; 0.9 the top 10%, and so on. In each case Leiden clustering was performed 40 times using the features obtained, with a different random seed each time. Shown in Fig. 4J-K are the mean and upper and lower quartiles of the enrichment scores obtained from these runs.

The outcomes support several of the previous conclusions. For example, in samples in which cells are numerous, sequencing is deep, and differences between cell types considerable, clustering succeeds regardless of how features are selected, as in the separation of CD8+ T cells from other T cells, Sod2+ macrophages from other macrophages, or FOXP3 T cells from other T cells, although in the last case, reducing the number of features too low by requiring too stringent a significance cutoff leads to a decrease in the reliability of clustering. In contrast, for the datasets with the smallest percentage of significant *ϕ*′ values, we find that restricting feature selection either by the relative magnitude of *ϕ*′, the statistical significance of *ϕ*′, or both, can improve performance.

### BigSur performs comparably to other methods, using fewer features

We compared BigSur to three of the most common methods for feature selection: HVGs (current default in scanpy), FindVariableFeatures (FvF, default in Seurat V4), and SCTransform (SCT) (Hafemeister & Satija, 2019; Hao et al., 2021; Wolf et al., 2018). HVGs ranks features by binning genes by normalized expression level, calculating the z-scores of genes with respect to their bin, and ranking the genes by their z-scores (Satija et al., 2015; Wolf et al., 2018). FvF fits the log of gene variance to gene mean and ranks genes by standardized residuals. (Hao et al., 2021). SCT fits each gene to a negative binomial model with sequencing depth as explanatory variable, then regularizes the parameters and ranks genes based on the resulting Pearson residual (Hafemeister & Satija, 2019).

We first compared the orders in which different methods ranked genes as features. We subsetted four datasets such that they only contained two groups of cells: a 1M PBMC dataset sequenced using Split-seq (containing CD4+ and CD8+ T cells); a keratinocyte dataset sequenced using 10x Genomics (basal and granular cells); and the 10k PBMC and macrophage datasets previously analyzed in Fig. 4 (CD4+ and CD8+ T cells, and M1 and M2 macrophages, respectively). Figure 5A shows Leiden clustering of these datasets using BigSur for feature selection (visualization of marker expression is also shown in Figure S4); similar clusters could be obtained using any of the other feature selection methods, with the exception of HVGs, which was unable to separate M1 and M2 macrophages (Figure S4).

**Figure 5:**
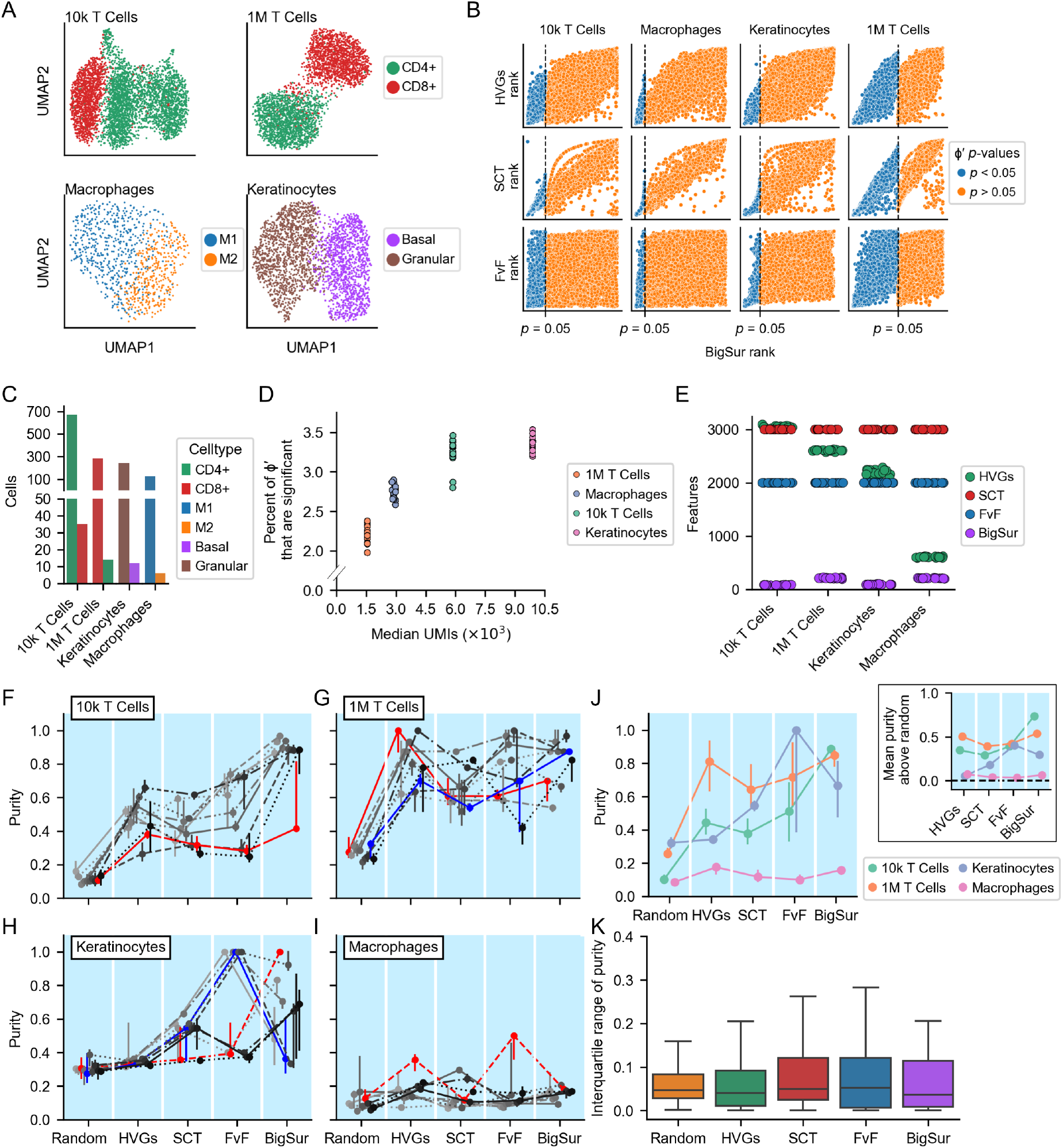
BigSur improves clustering performance using fewer total features. **A**. UMAPs of 4 datasets (two PBMC datasets, a keratinocyte dataset and a macrophage dataset; see main text) which were subset to contain only two biologically meaningful cell groups in each case. **B**. The ranking of features was calculated using HVGs, FvF, SCT and BigSur for each dataset (see main text for details). For BigSur, rankings are separately presented, in declining order of *ϕ*′, for genes with *p*-value below <0.05 (blue) and then for all other genes (orange); BigSur ranks are plotted against the ranks produced by other algorithms. **C**. For each dataset in A, we generated 20 “semi-synthetic” datasets comprising of a large population of cells and a rare population of cells. Bars show the cell numbers for each population (see methods; the individual steps are shown in Figure S5). **D**. The generation of such datasets was repeated 20 times and the percent of *ϕ*′ values that, in each case, displayed *p*<0.05 is plotted as a function of the median value of the total UMI per dataset. **E**. We selected features for each dataset in D using HVGs, SCT, FvF, and BigSur. The numbers of features were automatically selected using the first three methods’ defaults, and BigSur’s cutoffs were manually chosen in accordance with the results of panel D (see main text). **F–I**. For each of the semi-synthetic datasets in panel D, Leiden clustering, using 40 diferent random starting seeds, was performed using features that were either a random selection of 2000 genes, or those chosen by HVGs, SCT, FvF and BigSur. Purity scores were calculated (as in Fig. 2). Each line represents a different semi-synthetic dataset (for clarity, only 10 of the 20 are shown in each plot; see Fig. S5 for the remaining data), and the markers and bars are the medians and interquartile ranges (IQRs) of the purities (over the 40 different Leiden seeds). In panels F – I, the purities from selected semi-synthetic datasets are colored for discussion purposes (see main text). **J**. We calculated the medians of the purities of all the semi-synthetic datasets and grouped the medians by dataset. The markers represent medians (of the distribution of medians) and the bars represent IQRs. For the insert, we substracted the purities of each semi-synthetic dataset using random feature selection from the purities using the other methods. We then calculated the means of the resulting distribution (each marker represents a mean). **K**. Similar to J, we calculated the IQRs of the purities of all semi-synthetic datasets and grouped them by method.

We next compared how the rankings of genes as potential features by HVGs, FvF, and SCT, compared with the ranking (*ϕ*′) calculated by BigSur. Since BigSur both ranks genes and assigns *p*-values, we grouped genes into those that had FDR-adjusted *p*-values below 0.05, followed by those that did not. As shown in Fig. 5B, rankings produced by all three methods correlate to some degree with those produced by BigSur—with FvF showing the weakest and SCT the strongest correlation— but all of them highly ranked some genes that BigSur considered to be non-significant.

We also compared the speed of computation of each feature selection method. To do so, we randomly sampled increasing-size subsets of cells from the full 1M PBMC dataset and calculated the speed of each method for each set of cells (see Methods). BigSur was slower than HVGs and FvF but faster than SCT (Figure S4). At least 90% of the computation time for BigSur appears to be spent on *p*-value calculation (compare dashed line in Figure S4). Even with 100,000 cells, however, BigSur finishes in about a minute.

To create experimental data sufficiently challenging to discriminate between the performance of different methods, we downsampled the four datasets in Fig. 5A, randomly selecting a small population of cells from one of the two types and a larger population from the other type, so that the first type represented 5% of the total. The detailed steps in generating these “semi-synthetic” datasets are shown in Supplemental Figure 5 and described in Methods. Because sampling was random, the process was repeated 20 times, generating 20 independent datasets for each of the four cases.

The total numbers of cells in these semi-synthetic dataset cells ranged from 132 to 705 (Fig. 5C). When analyzed using BigSur, the percentages of *ϕ*′ values that were judged significant (*p* < 0.05) was relatively low in all cases, around 3%; median total UMIs varied between about 1,500 and 15,000 (Fig. 5D). Given the results in Fig. 4, we expected these conditions would make it relatively difficult to identify the rare cell type in these cases.

For each of the 80 semi-synthetic datasets, we ran the three common feature selection methods (using default parameters), as well as BigSur. In each case we also randomly chose a single set of 2000 genes to serve as random features.

The number of features selected by the different algorithms is shown in Fig. 5E. SCT and FvF uniformly chose 3000 and 2000 features, respectively. The number of features HVGs chose varied between a high of 3094 for the 10k T Cells semi-synthetic datasets and a low of 589 for the Macrophage semi-synthetic datasets. With BigSur, thresholds for feature selection were informed by the results in Fig. 4. Since the 10k T Cell and Keratinocyte semi-synthetic datasets displayed high median UMI/cell (around 5,870 and 9,900, respectively), we used a *ϕ*′ quantile cutoff of 0.99. Because the 1M T cell and macrophage semi-synthetic datasets both have low median UMI/cell (1561 and 2910 UMIs/cell, respectively) as well as a low numbers of cells (298 and 132), we used a more generous *ϕ*′ quantile cutoff of 0.9 (Fig. 4). In all, the number of features selected by BigSur ranged from 79 to 227, far fewer features than those selected by any other method.

We next used the selected features for clustering. As Leiden clustering can be sensitive to the randomly selected starting seed, we tested 40 independent starting seeds for each of the 80 datasets described above. We assessed performance in correctly isolating the rare cells, using the same purity score as in Fig. 2. For each of the four cases, we show results from 10 representative semi-synthetic datasets in Fig. 5F-5I, presenting the remaining 10 in Figure S5. Error bars illustrate the variation in results (interquartile range) introduced by the 40 random starting seeds. Fig. 5J summarizes the findings of Fig. 5F-5I. showing the median and interquartile range of purities obtained using each feature selection method.

For the 10k T Cells datasets (Fig. 5F), clustering using random features yielded purities ranging from 0.08 to 0.16. Using HVGs and SCT, purities ranged from 0.26 to 0.66 and 0.27 to 0.62, respectively. FvF yielded a wider range of purities, from 0.25 to 0.74. With BigSur, purities were higher than 0.8, with the exception of a single semi-synthetic dataset (colored in red, purity of 0.42), however this dataset was particularly sensitive to Leiden starting seed and for many starting seeds cases BigSur again produced very high purity. Overall, the average performance of features selected by BigSur was higher than with any other method (Fig. 5J).

For the 1M T Cells datasets (Fig. 5G), random feature selection yielded purities ranging from 0.22 to 0.33, whereas HVGs, SCT, FvF and BigSur produced similar maximal purities (1.0, 0.88, 1.0 and 0.97 respectively), albeit with considerable variability. Overall, BigSur produced the highest median purity (Fig. 5J).

With the Keratinocyte datasets, random feature selection and HVGs both performed poorly, yielding purities spanning 0.28 to 0.39 and 0.32 and 0.36 respectively (Fig. 5H). SCT, FvF and BigSur performance were all highly variable, varying from poor to very good depending on the dataset. The average performance of FvF exceeded that of BigSur (Fig. 5J), but the range was great for both methods.

Finally, with the Macrophage datasets, all methods yielded very poor purities, although an occasional dataset displayed slightly better purity when using either HVGs or FvF (Fig. 5I, colored in red).

Overall, even though no feature selection method consistently outperformed in all datasets, BigSur was the only method that had median scores above 0.6 for the first three datasets (Fig. 5J). While FvF, on average, outperformed BigSur on one dataset (Keratinocytes), the purity scores it produced were more widely spread than those of other methods, suggesting that the appropriateness of features selected by FvF is especially sensitive to the presence or absence of the individual cells in each dataset.

To assess whether sensitivity of clustering to Leiden starting seed varied systematically across different feature selection methods, we tabulated the sizes of the interquartile ranges for all the data in Fig. 5F-I and Fig. S5, and plotted the median and interquartile range of those ranges, as a function of selection method (Fig. 5K). Interestingly, BigSur had the smallest median value, followed by HVGs and random feature selection; and finally by SCT and FvF. A possible explanation for this effect is that the presence of false positive features increases the chance that the Leiden algorithm will be attracted to settle onto suboptimal solutions.

Taken together, the results argue that, despite using far fewer features, the performance of BigSur is generally as good as or better than other feature selection methods, and achieves greater reproducibility.

## DISCUSSION

Feature selection is an important general step in machine learning (Li et al., 2017), but the impact of different methods of feature selection on single cell transcriptomics has not been fully explored. Currently, several different methods are in common use as part of popular analysis packages, and multiple others have been proposed as improvements ((Andrews & Hemberg, 2018; Brennecke et al., 2013; Hafemeister & Satija, 2019; Jiang et al., 2016; Lall et al., 2022; Satija et al., 2015; Stuart et al., 2019; Su et al., 2021; Townes et al., 2019; Tyler et al., 2024)). Clear guidance about how to choose among them, or when to consider adjusting their parameters, is difficult to come by. Although some head-to-head comparisons have been published [e.g., (Germain et al., 2020; Su et al., 2021; Zhao et al., 2024), attention is not often paid to the difficulty of the task for which feature selection will be used. Here we focus on cell clustering and note that, in many cases—such as when cells are highly distinct in gene expression—feature selection hardly matters, and random sets of genes often do nearly as well as carefully chosen ones. Under these circumstances, large differences may exist in the accuracy with which different methods identify truly variable genes (e.g., (Andrews & Hemberg, 2018)) but they are likely to be of little practical importance.

In contrast, it is when using clustering to separate weakly dissimilar subsets of cells, especially when present as minor cell subpopulations, that feature selection method can really matter, especially when the total number of observations is relatively low. The impact of choice of feature selection method in this regime has been relatively unexplored, even though it may be expected to arise in scenarios of sequential subclustering of cell populations; characterizing cell heterogeneity in tumor samples; distinguishing intermediate states in cell lineages, and many other applications.

The results presented here argue that, in such cases, it can be important to use feature selection methods that not only identify genes that are truly variable but also eliminate ones that are not. Ultimately, the success of clustering must reflect a balance between signal (true positives) and noise (false positives). When the proportion of true positives is high, contamination by false positives may be irrelevant, but at lower signal strengths eliminating false positives is essential.

Here we develop a straightforward analytical approach, BigSur, that models the null distribution of gene expression observations as Poisson random variates from a distribution reflecting biological gene expression noise, the coefficient of variation of which is estimated from the data. For each gene, BigSur returns both a measure of variability *ϕ*′, and—by modeling the gene expression noise distribution as log-normal—the probability of observing that value by chance. The method automatically accounts for differences in data sparsity across genes, and differential sequencing depth across cells, so no data normalization or transformation is required. Whereas other feature selection methods sometimes also work by fitting data to distributions, those distributions are typically inferred from empirical observations of scRNAseq datasets, whereas with BigSur the use of the lognormal distribution to represent gene expression variability is grounded in both theory and observation (Bahar Halpern et al., 2015; Beal, 2017).

We compared the performance of BigSur against three popular selection methods, FvF (the default in Seurat), HVGs (the default in scanpy), and SCTransform (Hafemeister & Satija, 2019; Hao et al., 2021; Wolf et al., 2018). Both synthetic data, real data, and “semi-synthetic” datasets formed by guided downsampling of real data were used, concentrating on what we expected to be especially challenging regimes. Not only did these three methods highly-rank genes considered non-significant by BigSur, they tended to deliver much longer lists of features. For the most part, BigSur performed as well as or better than other algorithms (Fig. 5), despite using many fewer features, suggesting that, overall, it has a substantially better signal-to-noise profile. Consistent with this, results with BigSur also showed the lowest sensitivity to stochastic effects in Leiden clustering (Fig. 5K).

The fact that BigSur ranks features both by magnitude of variability and statistical significance gives investigators the opportunity to adjust cutoffs based on details of the data and the biological system it represents. Based on the observations in Fig. 4-5, we recommend routinely using an FDR-adjusted *p*-value threshold of 0.05, however the choice of 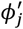 threshold should depend on the rarity of the cell type(s) one wishes to find (rare cells contribute less to 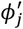) as well as whether cell-to-cell differences in gene expression are likely to be dominated by many small effects or a few large ones. In the absence of intuitions about these factors, we recommend starting with a quantile cutoff of 1%, increasing it as necessary to obtain at least a few hundred features. Observing the consistency of clustering performance over a range of quantile cutoffs may also be helpful, as may running the Leiden algorithm using more than one starting seed.

Here we did not assess the ability of BigSur to assist tasks other than cell clustering, such as pseudotime ordering, but suspect that the improvement in signal-to-noise ratio in feature selection will be helpful in that setting as well. Also, as noted above, the modified corrected Pearson residuals produced by BigSur may be used to generate modified corrected Pearson correlation coefficients, from which meaningful networks of gene expression correlation may be inferred, thanks to a dramatic reduction in false positive correlation (Silkwood et al., 2024).

## METHODS

### The Modified Corrected Fano Factor

The Fano Factor quantifies how much the variance of a distribution differs from that of a Poisson distribution. To the extent that scRNAseq data are not generally Poisson-distributed, the Fano factor does not reliably measure unexpected variability, but it can be modified, given an appropriate model, to do so. For any gene, assuming the null hypothesis (all cells draw from the same distribution), we model scRNAseq data as a random (Poisson) sample from a log-normal distribution, the mean of which is scaled in each cell by that cell’s sequencing depth.

In comparing data with the model, we avoid scaling individual observations by sequencing depth, as that distorts data distributions (Lun, 2018), and follow (Lause et al., 2021) in correcting Pearson residuals instead. The Pearson residual is a measure of each observation’s difference from the mean, scaled to the square root of the mean (expression 2, above). We correct it by using a different definition of mean in each cell, one that is scaled to account for total number of UMIs in the cell.

The corrected Pearson residual is then further modified to account for the expected greater variance of a compound Poisson log-normal distribution than a Poisson distribution. The variance of a compound distribution (assuming independence) is the sum of variances of the two underlying distributions. If the first is a Poisson Distribution, then *σ*^2^ = *μ*. For the second distribution, we equivalently express variance as *c*^2^*μ*^2^, the square of the coefficient of variation times the mean. As both distributions must share the same mean, the total variance is therefore *σ*^2^ = *μ* + *c*^2^*μ*^2^ = *μ*(1 + *c*^2^*μ*). We then use the square root of this expression to replace the denominator of the corrected Pearson residual. Introducing indices *i* and *j* to represent cell and gene, respectively, we obtain the following modified corrected Pearson residual 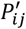:

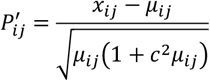

Just as the Fano factor, for gene *j*, may be expressed as an average over squared Pearson residuals, so may the modified corrected Fano factor be defined as

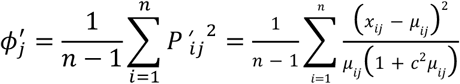

Where *n* is the number of cells. Note that the modified corrected Pearson residual can also be used to derive other useful statistics, such as a modified corrected Pearson correlation coefficient (Silkwood et al., 2024).

The analytical nature of the model underlying 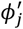 allows one to calculate, given a set of observations and cell sequencing depths, the probability that any given value of 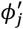 would arise by chance. Specifically, for each gene, we calculate the first 5 moments of 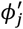 under the null hypothesis, from the moments of the individual Pearson residuals, which depend on the moments of the Poisson and log-normal distributions. Then we use numerical procedures (Cornish & Fisher, 1938) to estimate how far out an observed 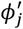 is on the tail of the null distribution, which may be expressed as a *p*-value. Derivation of the equations for moment calculation may be found in (Silkwood et al., 2024).

The only hyperparameter used in these calculations is *c*, which may be estimated from the behavior of an entire dataset, as described below. Note that the expectation value of 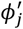 under the null hypothesis is unity regardless of the underlying distribution that describes gene expression. The choice of the log-normal distribution here only influences the expectation value for higher moments of *P*′, which are used in the calculation of *p*-values.

### Generation of Synthetic Datasets

To generate synthetic data, we first generate a log-normally distributed collection of mean expression values, i.e. a set of values < *y*_*j*_ >= 10^*x*^, where *x*∼*Normal*(*μ*_0_, *σ*_0_). In Fig. 3, we use *μ*_0_ = ™1 and *σ*_0_ = 0.8. Then, for each gene *j*, we generate a series of gene expression values as random variates of a log-normal distribution with mean < *y*_*j*_ > and coefficient of variation *c*. In Fig. 3, a value of *c* = 0.7 was used. Note that in many texts, log-normal distributions are parametrized in terms of the mean and standard deviation of the underlying normal distribution from which they may be derived, but here < *y*_*j*_ > and *c* refer to the actual mean and coefficient of variation of the log-normal distribution.

Given a matrix of synthetic gene expression values, for *n* cells and *m* genes:

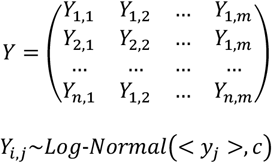

we replace each *Y*_*i,j*_ with a random variate from a Poisson distribution with rate parameter (mean) = *Y*_*i,j*_, to produce a final data matrix.

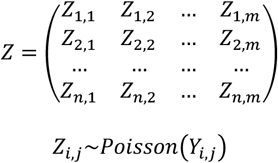

One could additionally simulate the effects of cell-specific variation in sequencing depth, by scaling each column in the Y matrix by an arbitrary constant prior to Poisson sampling, but this extra step was not carried out in Fig. 3.

The datasets in Fig. 3 simulate 2,000 cells expressing 15,000 genes each. The cells are divided into two equal sized groups of 1,000. For some of the genes (“not truly variable” genes), expression values for both groups are generated using a single < *y*_*j*_ >. For other genes (“truly variable” genes), the < *y*_*j*_ >used for the two groups of cells are independent, random selections from the set of < *y*_*j*_ >. Thus, the degree to which “truly variable” genes differ in expression between the two groups is itself distributed about a mean of zero.

### Fitting a coefficient of variation for underlying gene expression

Under the assumptions that *c* is approximately equal across genes, and that most of the genes in a dataset are not significantly differentially expressed across cells, one can estimate *c* by finding the value that minimizes the difference between 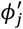 and 1, for the majority of genes. Because *c* influences the modified corrected Pearson residual only through the term 1 + *c*^2^*μ*_*ij*_, it follows that the choice of *c* has little influence on 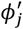 when *μ*_*ij*_<<1. Typically we learn *c* by finding the value that minimizes the absolute value of the slope of a linear fit to a plot of 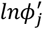 vs. *lnμ*_*j*_, for *μ*_*j*_ ∈ [0.01,10].

### Dimensionality reduction and clustering

Dimensionality reduction and clustering were done using scanpy v1.8.2. The PBMC 10k dataset was filtered to retain only genes with expression in at least 3 cells and cells with at least 400 expressed genes and no more than 10% of UMI coming from mitochondrially-encoded genes. The retina (Menon et al., 2019), keratinocyte (Guerrero-Juarez et al., 2022), 1 M PBMC dataset and macrophage (Carvalho et al., 2021) datasets were filtered similarly, except cells were not subsetted based on their mitochondrial gene content. Counts were normalized as follows:

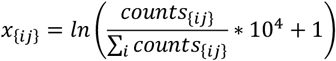

Where *x*_{*ij*}_ is the normalized count of gene *j* in cell *i*.

During subsequent subsetting, we removed genes that were expressed in fewer than 10 cells. Principal component analysis was performed on the normalized data and the top 50 principal components (PCs) were retained. The *k*-nearest neighbors graph was calculated from the PCs, and UMAP dimensions calculated using scanpy default parameters. Clusters were identified using the Leiden algorithm implemented by scanpy with default parameters and, if not otherwise stated, resolution = 1 and random seed = 0 (default).

### Classification of cell types using random genes

Cell types in the 10k PBMC dataset were identified using marker genes on clusters calculated as specified above, using HVGs with defaults as feature selection. scrublet v0.2.3 (Wolock et al., 2019) was used to identify doublets using default parameters. For every set of randomly selected genes (uniform sampling using numpy.default_rng.choice function, setting replace = False), PCA was calculated using sklearn’s implementation of TruncatedSVD with number of PCs = 50, except for 25 genes where number of PCs = 25. A linear support vector machine (SVM, using sklearn.svm.SVC) was trained on each set of PCs calculated by TruncatedSVD, with ground truth labels being the cell types identified as described above. The adjusted Rand index (ARI) and normalized mutual information (NMI) scores were calculated by comparing predicted labels of the SVM to ground truth labels, using sklearn’s implementation of both scores.

### Generation of semi-synthetic datasets

To compare feature selection methods, we devised a procedure that creates datasets of modest size, in which there are two populations of relatively similar but transcriptionally distinct cells, one substantially rarer than the other. The procedure entails taking pairs of related cells (e.g. CD4+ and CD8+ T cells, basal and granular keratinoctyes, etc.) that had previously been definitively identified, and mixing small numbers of them together.

Briefly, we first clustered the data shown in Fig. 5A, extracted the desired cell pairs, and reclustered to ensure the identification of two homogeneous groups. Then we combined all of the cells from the larger of the two groups with a random subset of cells from the other, of sufficient size so that the resulting ratio of cell types was 19:1 (the absolute cell numbers are reported in Fig. 5C). We then removed genes expressed in fewer than three cells. The cell barcodes of each semi-synthetic dataset were also saved (see data availability).

### Purity score

For any group of target cells that we subject to clustering, we may define a purity score for those cells in a round of clustering as the maximum fraction of each cluster obtained that consists of target cells.

When discussing the purity score for synthetic data in which there are two pre-determined groups of cells (Fig. 3B, 3F), we chose to present the mean of the purity scores calculated for each of the two cell types. Clusters were generated from the data using a resolution parameter of 0.1. In Fig. 3B, ten rounds of random feature selection and Leiden clustering were performed on each dataset and the average results across the ten trials is presented. In Fig. 3F, Leiden clustering was performed 50 times using unique random seeds and results were averaged across all trials.

### Speed comparisons

To compare the speed of HVGs, SCT, FvF and BigSur, we used the 1M PBMC dataset from Parse Biosciences. The dataset was filtered to retain only cells with more than 400 genes and genes expressed in at least 3 cells. For the comparison between HVGs and BigSur, 10 datasets with increasing numbers of cells (100, 500, 10^3^, 5*10^3^, 10^4^, 5*10^4^, 10^5^ and 2.5*10^5^ cells) were randomly sampled from the 1M PBMC dataset. Each sampled dataset was filtered to retain only genes expressed in at least 1 cell.

Since SCT and FvF were first implemented in R, we randomly sampled a 2.5*10^5^ dataset from the filtered dataset and exported to R. 10 datasets were randomly sampled from the filtered dataset per number of cells, and the features were calculated from the sampled datasets. All real dataset results were generated on an iMac Pro (2017) with 10 3 Ghz Intel Xeon W cores and 256 GB of RAM running Ventura 13.2.1.

## Data availability

No new data were generated as part of this study. The 10k PBMC dataset was downloaded from 10x Genomics, https://cf.10xgenomics.com/samples/cell-exp/6.1.0/10k_PBMC_3p_nextgem_Chromium_X/10k_PBMC_3p_nextgem_Chromium_X_raw_feature_bc_matrix.h5. The 1 million PBMC dataset was downloaded from Parse Biosciences, https://cdn.parsebiosciences.com/1M_PBMC_T1D_Parse.zip. The retina (Menon et al., 2019), macrophage (Carvalho et al., 2021) and keratinocyte (Guerrero-Juarez et al., 2022) datasets were downloaded from GEO, (GSE137537, GSE164498 and GSE141526 respectively). The metadata for each dataset, including the cell barcodes of each semi-synthetic dataset, are included in supplementary data.

## Code availability

Code is freely available as a python package on github: https://github.com/landerlabcode/BigSur.

An R version can be found at: https://github.com/landerlabcode/BigSurR.

A Mathematica version can be found at: https://github.com/landerlabcode/BigSurM.

## ACKNOWLEDGEMENTS

The authors thank Fairlie Reese, Samuel Morabito and Tatyana Lev for assistance with code optimization, and Joshua Gervin for helpful discussions during preparation of the manuscript. Research support was provided by NIH grants U54CA217378, P30AR075047, R01DE019638, R01AR081671, R01GM152494 and R01CA237563. ED and KS were also supported by training grant T32GM136624.

