## Supplementary Figures for "Statistically principled feature selection for single cell transcriptomics"

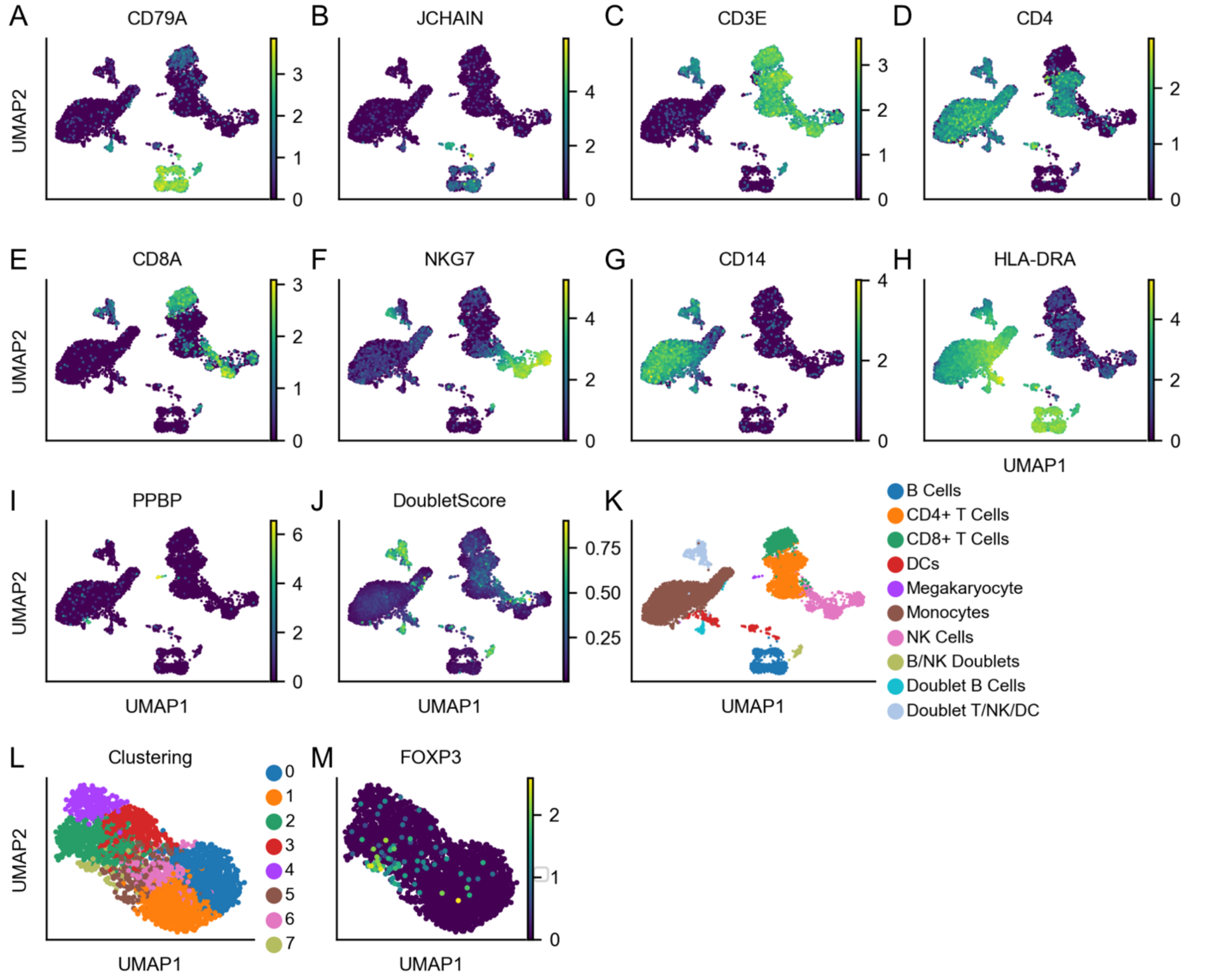


**Supplementary Figure 1: Celltype identification in the PBMC dataset.** Features for the 10k PBMC dataset were selected using HVGs defaults. Clusters were assigned using the Leiden algorithm. **A – B.** *CD79A* and *JCHAIN* were used to identify B cells. **C – E.** Markers to identify the T cells: *CD3E* for all T cells, *CD4* for CD4+ T cells, *CD8A* for CD8+ T cells. **F.** *NKG7* was used to identify NK cells. **G.** *CD14* was used to identify monocytes. **H.** Co-expression of *HLA-DRA* and *CD14* was used to identify dendritic cells (DCs). **I.** *PPBP* was used to identify Megakaryoctes. **J.** Scrublet was used to calculate the doublet score for each cell (see Methods). **K.** Celltypes identified using the above markers; same panel as in Figure 1A. **L.** The CD4+ T cells were subset. Features of this subset were selected using HVGs, and dimensionality reduction and clustering were performed as above. **M.** *FOXP3* was used to identify Tregs.

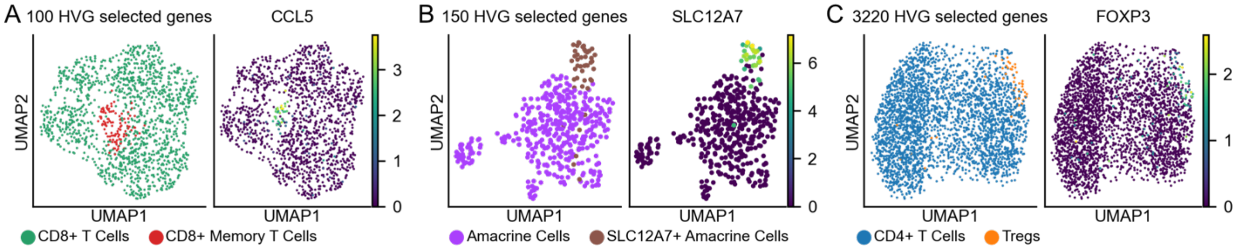


**Supplementary Figure 2: Identification of cell groups used in Figure 2. A.** CD8+ T cells were subset from the 10k PBMC dataset. The 100 top HVG features were selected, and dimensionality reduction and clustering were done with defaults. The CD8+ memory T cell population was identified by expression of *CCL5*. **B.** Amacrine cells were subset from the retina dataset and reclustered using 150 top HVG. We identified a cluster of *SLC12A7*-expressing amacrine cells. **C.** The CD4+ T cells were subset from the 10k PBMC dataset and re-clustered using the top 3220 HVG features. Tregs were identified using *FOXP3* expression.

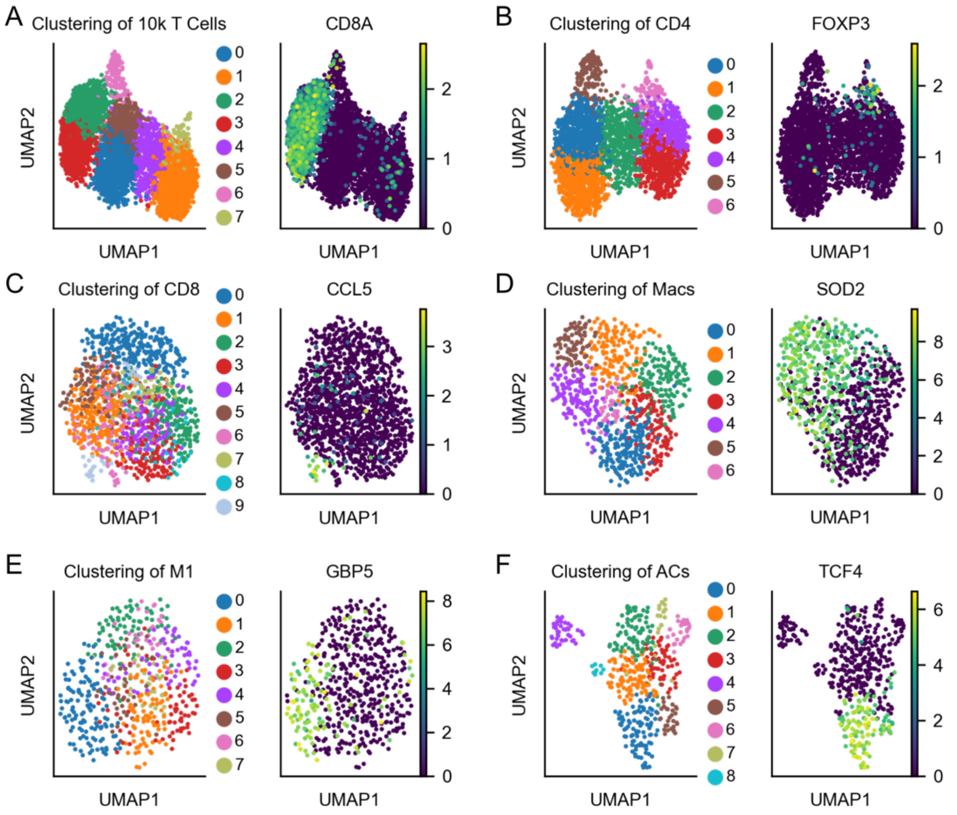


**Supplementary Figure 3. A – F.** Markers and clusters for each of the datasets used for the enrichment calculations in Figure 4J and K. Features were ranked using BigSur. Dimensionality reduction was performed using features with *p*-values smaller than 0.05 and $\phi'$ quantile cutoffs of 0.9 for the 10k T Cells, CD4 T Cells, Macs, and ACs datasets and 0.99 for the CD8 T Cells and M1 macrophages.

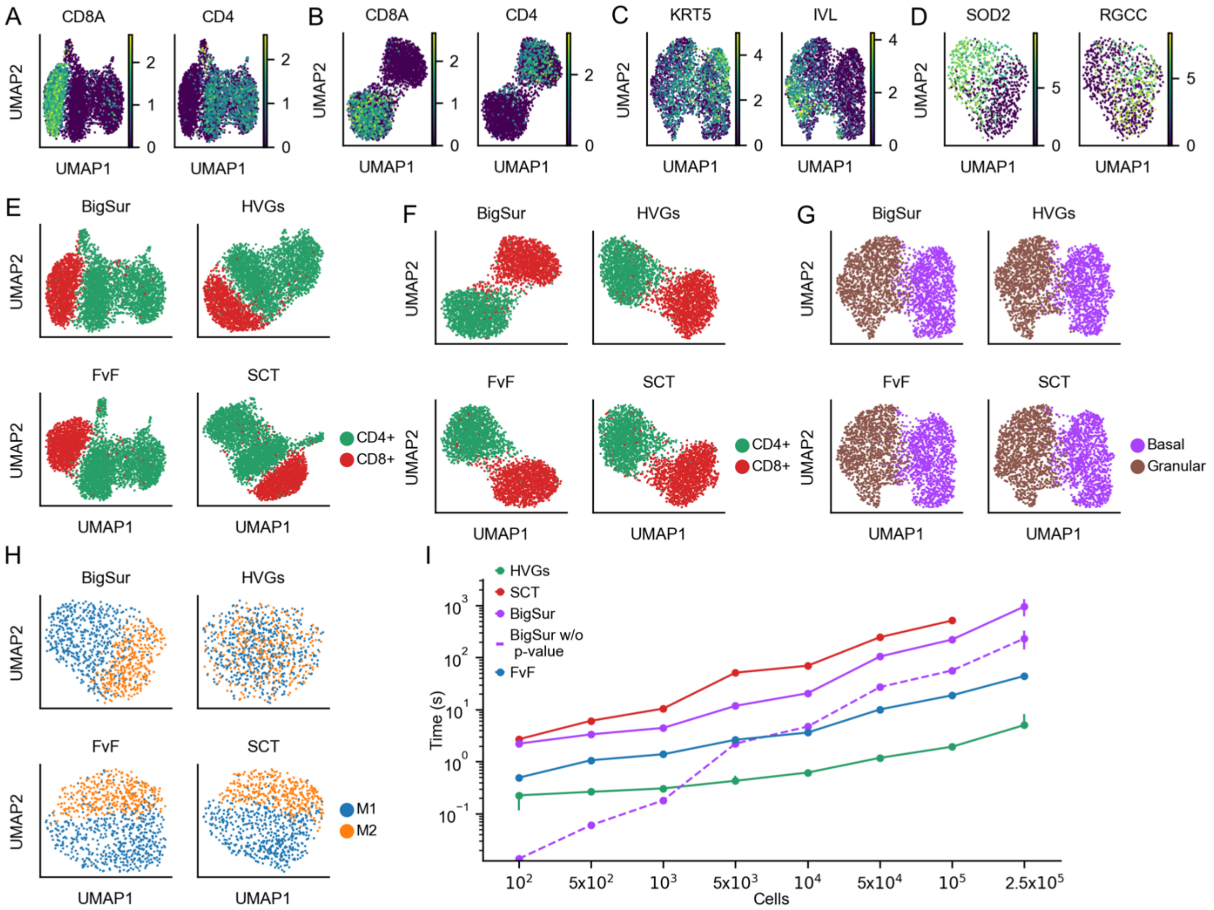


**Supplementary Figure 4. A – D**. Markers for each of the groups of cells for each of the datasets shown in Figure 5A: 10k T Cells (panel A), 1M T Cells (panel B), Keratinocytes (panel C) and Macrophages (panel D). **E – H.** Features for each dataset in Figure 5A (10k T Cells, 1M T Cells, Keratinocytes and Macrophages for panels E, F, G and H respectively) were selected using each of the four feature selection methods and clustering was performed. **I.** The computational run time of each method on different dataset sizes was recorded (see Methods). Markers are medians and bars are interquartile ranges (IQRs). We were unable to run SCT on datasets with $2.5 \times 10^5$ cells due to lack of memory.

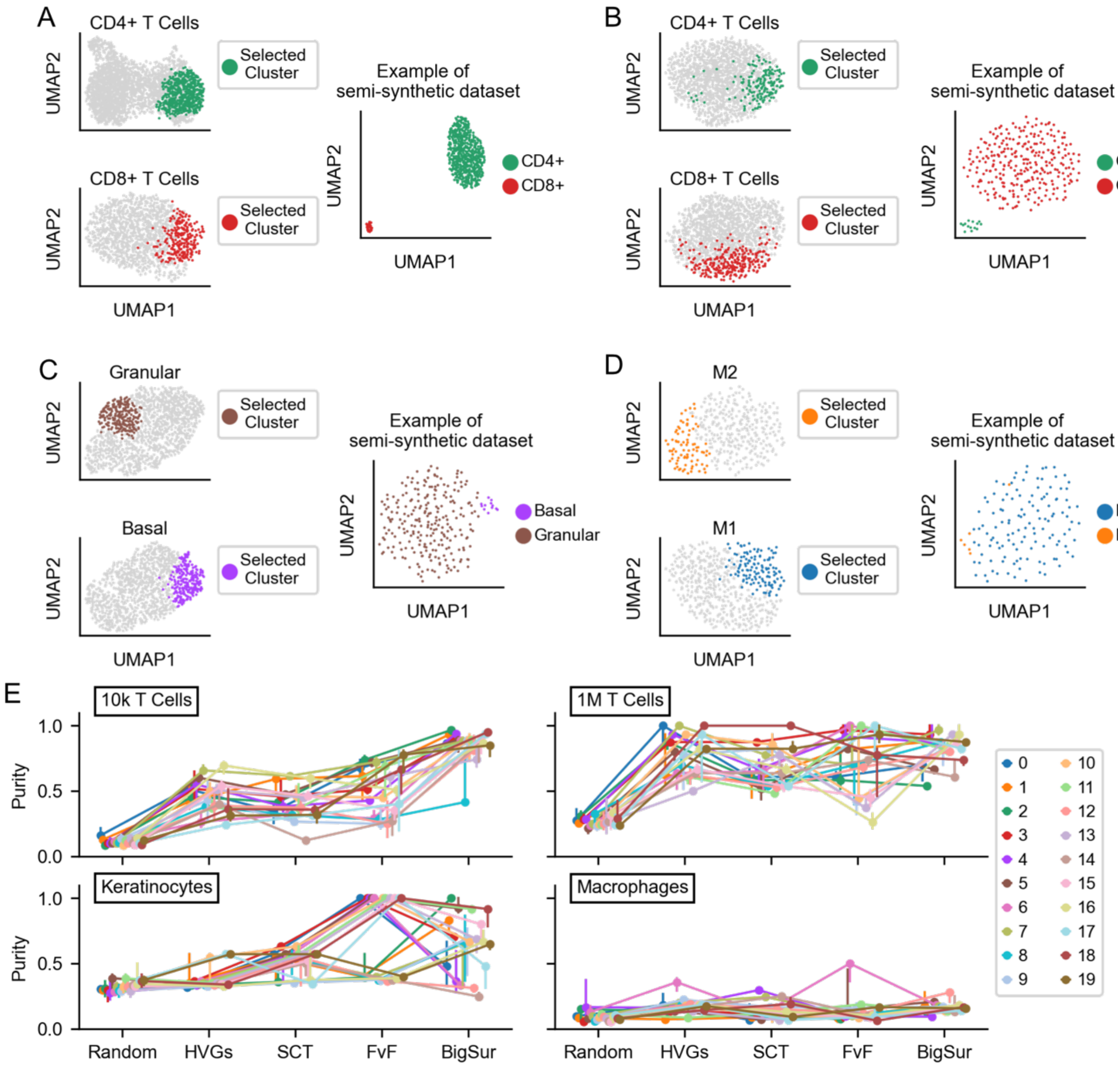


**Supplementary Figure 5. A – D.** Visualization of intermediate steps (described in Methods) for semi-synthetic data generation, along with an example of a semi-synthetic dataset, for the 10k T Cells (panel A), 1M T Cells (panel B), Keratinocytes (panel C), and Macrophages (panel D) datasets. **E.** The purities of all semi-synthetic datasets using each feature selection method. The first 10 datasets (0-9) are the datasets shown in Figure 5F – G.
